# The cancer chemotherapeutic 5-fluorouracil is a potent inhibitor of *Fusobacterium nucleatum* and its activity is modified by the intratumoral microbiota

**DOI:** 10.1101/2022.03.12.484096

**Authors:** Kaitlyn D. LaCourse, Alexander Baryiames, Andrew G. Kempchinsky, Christopher D. Johnston, Susan Bullman

## Abstract

*Fusobacterium nucleatum* is among the most prevalent and dominant bacterial species in colorectal cancer (CRC) tumor tissue, and growing evidence supports its role in cancer progression and poorer patient prognosis. Here we perform a small molecule inhibitor screen of 1,846 bioactive compounds against a CRC isolate of *F. nucleatum* and find that 15% of inhibitors are antineoplastic agents including fluoropyrimidines. Validation of these findings reveal that 5-fluorouracil (5-FU), the first-line chemotherapeutic used to treat CRC worldwide, is a potent inhibitor of *F. nucleatum* CRC isolates at concentrations found in serum of CRC patients treated with 5-FU. We also identify members of the intratumoral microbiota that are resistant to 5-FU, including *Escherichia coli*. Further, CRC *E. coli* isolates can modify 5-FU and relieve 5-FU toxicity towards otherwise-sensitive *F. nucleatum* and human CRC epithelial cells. Lastly, we demonstrate that *ex-vivo* CRC tumor microbiota from patients undergo different levels of community disruption after 5-FU exposure and have the potential to deplete 5-FU levels, thereby reducing local drug efficacy. Together, these observations argue for further investigation into the role that the CRC intratumoral microbiota plays in patient response to chemotherapy.

## Introduction

Colorectal cancer (CRC) tumor cells exist within a complex microenvironment that include intimate associations with certain members of the bacterial microbiota^1^. Worldwide, genomic analyses consistently reveal enrichment of the bacterial species *Fusobacterium nucleatum* (*Fn*) in human colon cancers relative to noncancerous colon tissues^2–8^. Several studies over the last decade have revealed that *Fn* contributes to tumorigenesis and accelerated cancer cell growth^9–14^. A high burden of *Fn* in the tumors of CRC patients is associated with resistance to chemotherapy, disease recurrence, metastasis, and poorer survival^6,7,14–18^. The benefit of chemotherapy is limited for CRC patients^19^, especially in late-stage disease, highlighting the clinical need for more effective treatments to combat this disease. Our prior research demonstrated that treatment of *Fn*-positive human CRC xenografts in mice with the antibiotic metronidazole significantly reduced tumor growth and cancer cell proliferation^18^, suggesting that targeting *F. nucleatum* could be a novel therapeutic approach for a subset of CRC patients. Owing to the broad-spectrum antimicrobial activity of metronidazole, which can also target beneficial members of the gut microbiota, an initial focus of this work was to identify *F. nucleatum* inhibitors with narrow spectrum activity via a small molecule screen of 1,846 bioactive compounds.

Surprisingly, our screen identified 5-fluorouracil (5-FU), the primary chemotherapeutic used to treat patients with CRC, as a potent inhibitor of *Fn* CRC strain growth. By exploring 5-FU toxicity towards other dominant CRC-associated bacterial species that routinely co-occur with *Fn*, we show that *Escherichia coli, Bacteroides fragilis, Bifidobacterium breve*, and *Parvimonas micra* isolates from CRC tumors are resistant to physiologically relevant concentrations of 5-FU. Analysis of ex-*vivo* CRC microbiota in the presence of 5-FU demonstrates that bacterial community members can resist 5-FU toxicity and potentially reduce drug bioavailability, thereby protecting both CRC tumor cells and sensitive bacterial strains. These data suggest that the interplay between the intratumoral microbiota and chemotherapeutics could inform the design of individual CRC patient treatment regimes.

## Results

### Bioactive compound library screening for inhibitors of *Fusobacterium nucleatum*

Initially, we sought to determine which classes of drugs could inhibit *F. nucleatum* (*Fn*) growth. In a pilot assay, we screened a random selection of 1,846 small molecules [32 μM] from the Broad Institute’s “Bioactive Compound” library for their ability to inhibit the growth of a human CRC isolate *Fn* subsp. *animalis* (*Fna*) SB010^18^ (Figure 1A and Table S1). We identified 34 inhibitory compounds of *Fn* growth in broth culture (Figure 1B and Figure S1A). Approximately half of those identified (56%) are known antimicrobial compounds. Interestingly, 15% of inhibitors are classified as antineoplastic agents that act upon either thymidylate synthase, estrogen receptors, or topoisomerase II (Figure 1C). Among these, the mainstay CRC chemotherapeutic drug Tegafur and its active metabolite 5-fluorouracil (5-FU) were identified.

**Figure 1:**
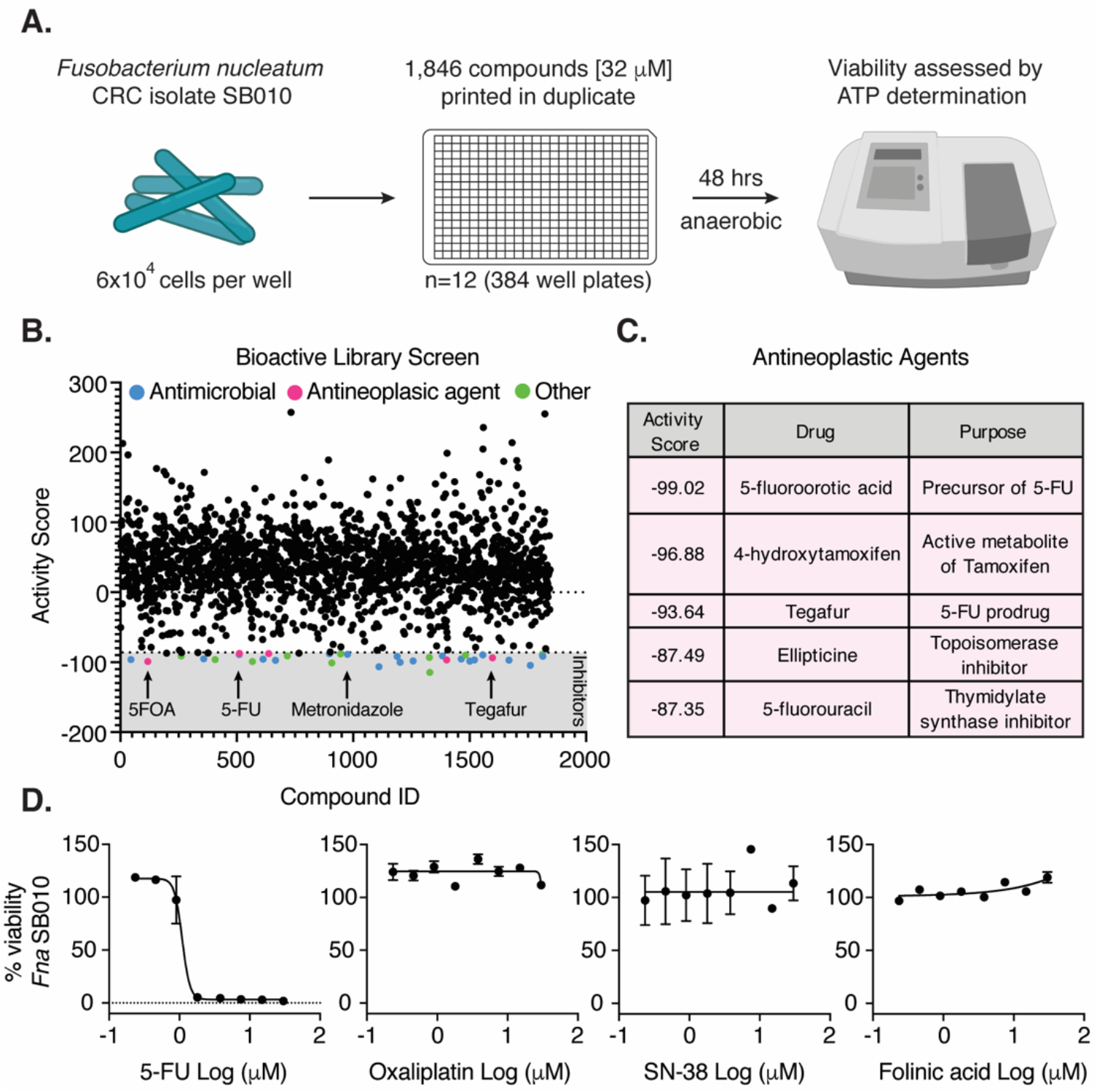
A bioactive library screen identifies several antineoplastic agents as *Fusobacterium nucleatum* growth inhibitors. (A) Schematic describing the small molecule screen workflow to determine *F. nucleatum* drug sensitivity (B) Activity scores for n=1,846 compounds on *F. nucleatum* viability, determined through ATP measurements. Inhibitors are compounds that are three standard deviations away from the neutral control (media alone). (C) A list of the antineoplastic agents found to be *F. nucleatum* inhibitors arranged by their respective activity scores. (D) Eight-point dose-response curves of *F. nucleatum* viability after exposure to 0.25-32 μM of the active metabolites for various chemotherapeutics used to treat metastatic colon cancer for 48 hours. The connecting line is a non-linear regression of the log(inhibitor) vs. response with a variable slope (four-parameters).

To validate and expand upon these initial findings, we monitored *Fn* growth in the presence of 24 chemotherapeutics using 8-point dose-response curves ranging from 0.23-30 μM. We found 5-FU and its prodrugs Tegafur and Carmofur were potent inhibitors of *Fn* growth (IC50: 0.46 and 29.98 μM respectively) (Figure 1D, Figure S1C-D, and Table S2). Capecitabine, another prodrug of 5-FU, was found to have no impact on *Fn* growth under these conditions, possibly due to the inability of *Fn* alone to metabolize the drug to its active form (Figure S1E). In clinical settings, metastatic CRC patients are treated with 5-FU stabilized with folinic acid in combination with either oxaliplatin or SN-38, the active metabolite of irinotecan (FOLFOX or FOLFIRI respectively)^20^. We found that these compounds did not significantly impact *Fn* growth *in vitro* (Figure 1D). These initial data suggest that standard chemotherapy for CRC could concomitantly inhibit *Fn* growth at the tumor site.

### 5-fluorouracil is a potent inhibitor of *F. nucleatum* CRC clinical isolates

*Fn* is one of the most dominant CRC-associated bacterial species^2,3^; however, sensitivity of CRC *Fn* clinical isolates to 5-FU, the primary chemotherapeutic used to treat CRC, has not been previously explored. We therefore evaluated the half-maximal inhibitory concentration (IC50) of 5-FU in 14 strains of *Fn*, comprising isolates from CRC tumor tissue (n=11), the oral cavity (n=2), and inflamed irritable bowel disease tissue (n=1). IC50 values were determined from 8-point dose-response curves generated after 48 hours of incubation with 5-FU, a time frame chosen to reflect a typical two-day continuous infusion of 5-FU within a clinic setting. We defined *sensitivity* as IC50 values lower than the concentration of 5-FU found in patient sera (2.5-10 μM)^21–23^, and *resistance* as IC50 values higher than this range. These analyses revealed that 5-FU is a potent inhibitor of the majority of *Fn* strains tested with IC50 values ranging from 0.14-4.3 μM (Figure 2A and Figure S2A-J). Genomic variability within *Fn* is extensive and currently each strain can be assigned into one of four subspecies; subsp. *nucleatum* (*Fnn*), *animalis* (*Fna*), *polymorphum* (*Fnp*), *vincentii/fusiforme* (*Fnv*)^24,25^. *Fna* is the most prevalent subspecies found in CRC in epidemiological studies^26,27^, and we therefore sampled *Fna* at a higher frequency (10/14 isolates tested). We confirmed that 5-FU is a potent growth inhibitor of CRC *Fn* isolates representing all four subspecies, suggesting that 5-FU sensitivity is a core feature of *Fn* species (Figure 2B). These findings indicate that the growth of *Fn* is inhibited by 5-FU exposure at physiologically relevant levels.

**Figure 2:**
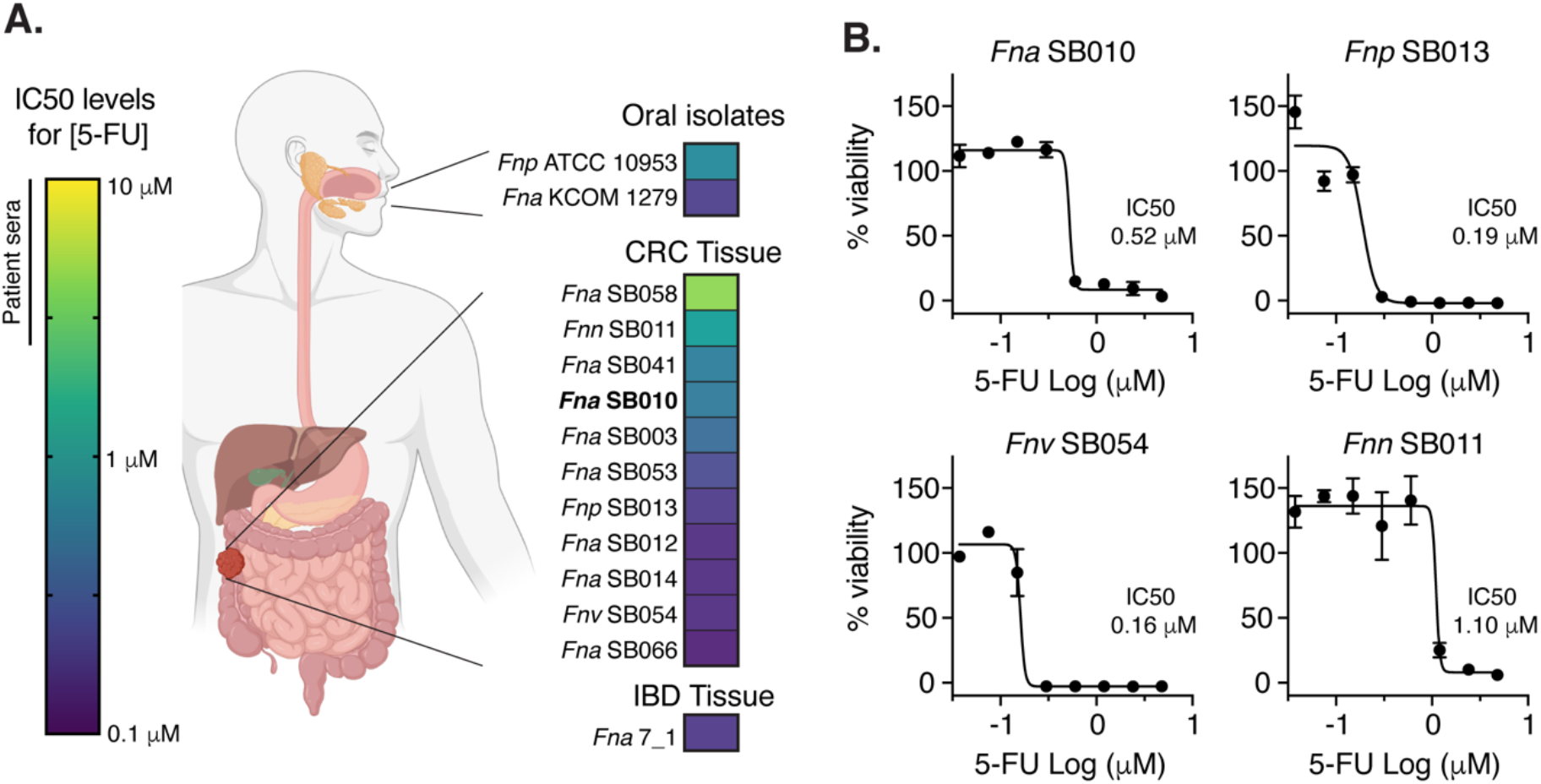
*F. nucleatum* isolates are sensitive to physiological concentrations of 5-fluorouracil. (A) Heat map depicting the concentrations of 5-FU where *F. nucleatum* isolates (n=14) are 50% viable. The clinical isolates are grouped by the sampling source. The level of 5-FU in patient sera is labelled between 2.5-10 μM. (B) Eight-point dose response curves of *F. nucleatum* viability for a single representative of each subspecies: *animalis, polymorphum, vincentii*, and *nucleatum* after exposure to 0.0375-4.8 μM 5-FU for 48 hrs. The connecting line is a non-linear regression of the log(inhibitor) vs. response with a variable slope (four-parameters).

### A CRC isolate of *Escherichia coli* modifies 5-fluorouracil and abrogates toxicity toward *F. nucleatum* and CRC epithelial cells

Post-chemotherapeutic treatment, studies have found *Fn* persists in distant CRC metastases^18^, and a high load of *Fn* in primary CRC tissue is positively correlated with increased risk of disease re-occurrence^14,16^. Furthermore, in patients with locally advanced rectal cancer, *Fn* positivity after 5-FU-based neoadjuvant chemotherapy significantly increases the risk of relapse^16^. This suggests that in a subset of patients *Fn* can endure chemotherapy despite 5-FU toxicity and promote disease progression in patients harboring these strains. We hypothesized that co-occurring bacterial species in CRC tumors might protect *Fn* from 5-FU through sequestration or modification of the compound, lowering drug toxicity toward nearby cells in colonized microniches.

5-FU is a uracil analog that acts as a pan-antimetabolite and inhibits the activity of thymidylate synthases, enzymes that are highly conserved between humans and bacteria^28^. To test whether 5-FU acts as a broad antimicrobial toward bacterial species commonly found in CRC, we measured 5-FU sensitivity of CRC tumor isolates of *Bacteroides fragilis, Escherichia coli, Bifidobacterium breve*, and *Parvimonas micra* using 13-point dose-response curves. All species tested were resistant to physiologically relevant concentrations of 5-FU (2.5-10 μM) (Figure 3A). Therefore, we reasoned that these strains may harbor mechanisms capable of detoxifying 5-FU. We monitored the concentration of extracellular 5-FU in the presence of these four bacterial strains over 48 hours using liquid chromatography-mass spectrometry (LC-MS) and observed 5-FU depletion from extracellular media specifically in the presence of the CRC *E. coli* strain within 24 hours (Figure 3B). Local depletion of 5-FU has the potential to relieve toxicity on nearby sensitive bacterial species. To test if bacterially mediated 5-FU depletion influences *Fn* viability, we cultured *Fna* SB010 in conditioned media with 4 μM 5-FU (~8 times the IC50 of this strain) that was previously exposed to *E. coli* or *B. fragilis* cells. Only *E. coli* modified 5-FU rescued *Fna* SB010 growth. *B. fragilis*, which did not to deplete 5-FU from media, failed to abrogate the drugs toxicity (Figure 3C). Additionally, we assessed a colibactin positive (*pks*+) strain of *E. coli*, owing to their prevalence in up to 55% of CRC patient tissues ^29,30^, and similarly observed rescue of *Fna* SB010 growth (Figure 3C). Addition of fresh 5-FU to these bacterial supernatants led to inhibition of *Fna* SB010 suggesting that *Fn* protection against 5-FU was mediated directly by *E. coli* cells and not due to an unknown protective component present in the supernatant (Figure 3C). Furthermore, inoculation of *E. coli* and *Fna* SB010 together at T0 of 5-FU exposure protects *Fna* SB010 from 5-FU toxicity (74% viability after 48 hours; p=0.0002) (Figure S3A). LC-MS confirmed that 5-FU disappearance occurs within 8-24 hours in multispecies co-cultures of *Fna* SB010, *E. coli*, and *B. fragilis* (Figure S3B).

**Figure 3:**
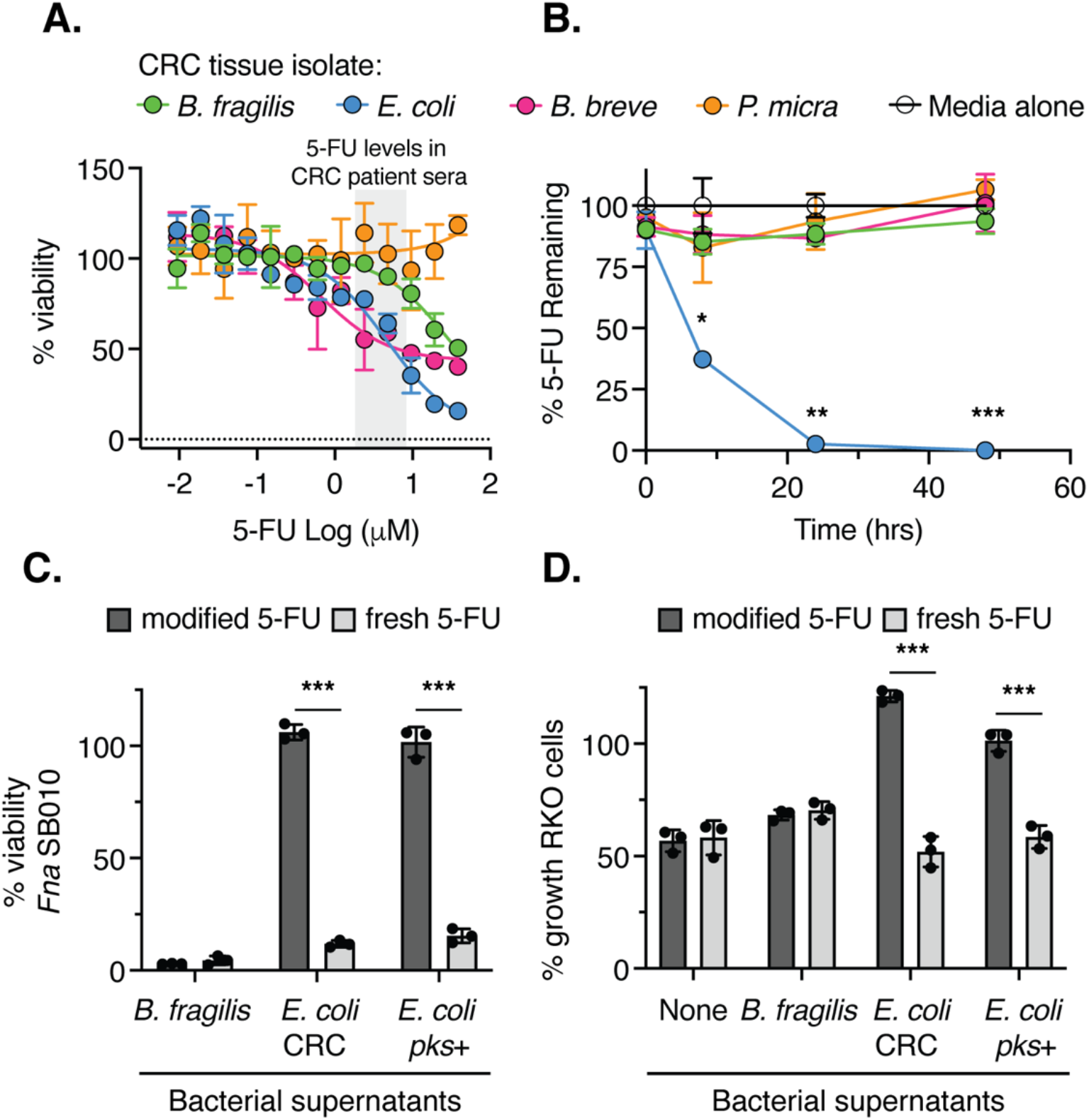
*E. coli* isolates deplete 5-fluorouracil and reduces drug toxicity toward *F. nucleatum* and human CRC cells. (A) Thirteen-point dose-response curves of *B. fragilis, E. coli, B. breve*, and *P. micra* viability after exposure to 0.0023-38.4 μM 5-FU for 48 hours. The range of 5-FU in patient sera is labelled between 2.5-10 μM. The connecting lines are a non-linear regression of the log (inhibitor) vs. response with a variable slope (four-parameters). (B) Measurement of 5-FU (4μM) disappearance in the supernatant when exposed to the indicated bacterial strains or media alone for 0, 8, 24, and 48 hours. 100% is set to the media-alone condition at 0 hrs. (C) The viability of *Fna* SB010 when incubated in the conditioned supernatants of the indicated bacterial species for 48 hours. The conditioned supernatants of the indicated bacterial species were incubated with 5-FU (4 μM) for 48 hours (dark grey bars) or had fresh 5-FU (4 μM) added immediately prior to sterilization using a 0.2 μm filter (light grey bars). 100% is set to the *F. nucleatum* viability in the indicated bacterial supernatants alone for 48 hrs. (D) The relative growth of RKO CRC epithelial cells when incubated in the conditioned supernatants of the indicated bacterial species for 72 hours. The conditioned supernatants of the indicated bacterial species were incubated with 5-FU (20 μM) for 48 hours (dark grey bars) or had fresh 5-FU (20 μM) added immediately prior to sterilization using a 0.2 μm filter (light grey bars). This supernatant was diluted ¼ when added to the RKO culture, resulting in anticipated 5 μM 5-FU. 0% growth is the confluency of the RKO cells in each condition at 0 hours. 100% growth is the confluency of the RKO cells with incubated with the indicated bacterial supernatant alone for 72 hours. For B-D: *indicates p-values <0.05, ** indicates p-values <0.01, and ** indicates p-values <0.001 as determined by a Student’s t-test.

These findings led us to question if *E. coli* modification of 5-FU could alter its chemotherapeutic efficacy toward CRC cells. To evaluate this, we cultured RKO cells, a human CRC epithelial cell line reported to be sensitive to 5-FU (IC50: 5 μM)^31^, with bacterially modified 5-FU (5 μM) and observed cell growth over 72 hours. Remarkably, prior exposure to *E. coli* (CRC and *pks*+ strains) completely abrogated 5-FU toxicity against human CRC epithelial cells (Figure 3D). In total, these results suggest that bacteria-mediated depletion of 5-FU reduces drug efficacy against neighboring bacteria cells and human CRC tumor epithelial cells.

### CRC patient-derived *ex-vivo* tumor microbiota interacts with 5-fluorouracil

Based our findings that certain members of the microbiota have the ability to detoxify 5-FU, we next sought to determine if 5-FU exposure affects the CRC microbiota community structure by facilitating expansion of 5-FU resistant bacterial species. To generate ex-*vivo* CRC communities, treatment naïve CRC tissue (n=6 patients) was manually homogenized in bacterial broth and passaged through a 16-gauge needle until homogenous. Human cells and debris were pelleted by gentle centrifugation at 300×g leaving the tissue-associated microbiota in suspension. The remaining bacterial cells were incubated either in broth only or in the presence of a high dose of 5-FU (30 μM) under anaerobic conditions for 48 hours. The resulting community structure was then determined through metagenomic sequencing (Figure 4A, Table S2, and Table S3). Bacterial presence in these patient tumors was examined prior to dissociation via RNAscope fluorescence in situ hybridization (FISH) (Figure 4B and Figure S4A-F). Metagenomic analyses revealed that indeed exposure to 5-FU alters the relative abundance of community members, allowing for the expansion of 5-FU resistant bacterial species (patients CRC_01, CRC_05, CRC_06).

**Figure 4:**
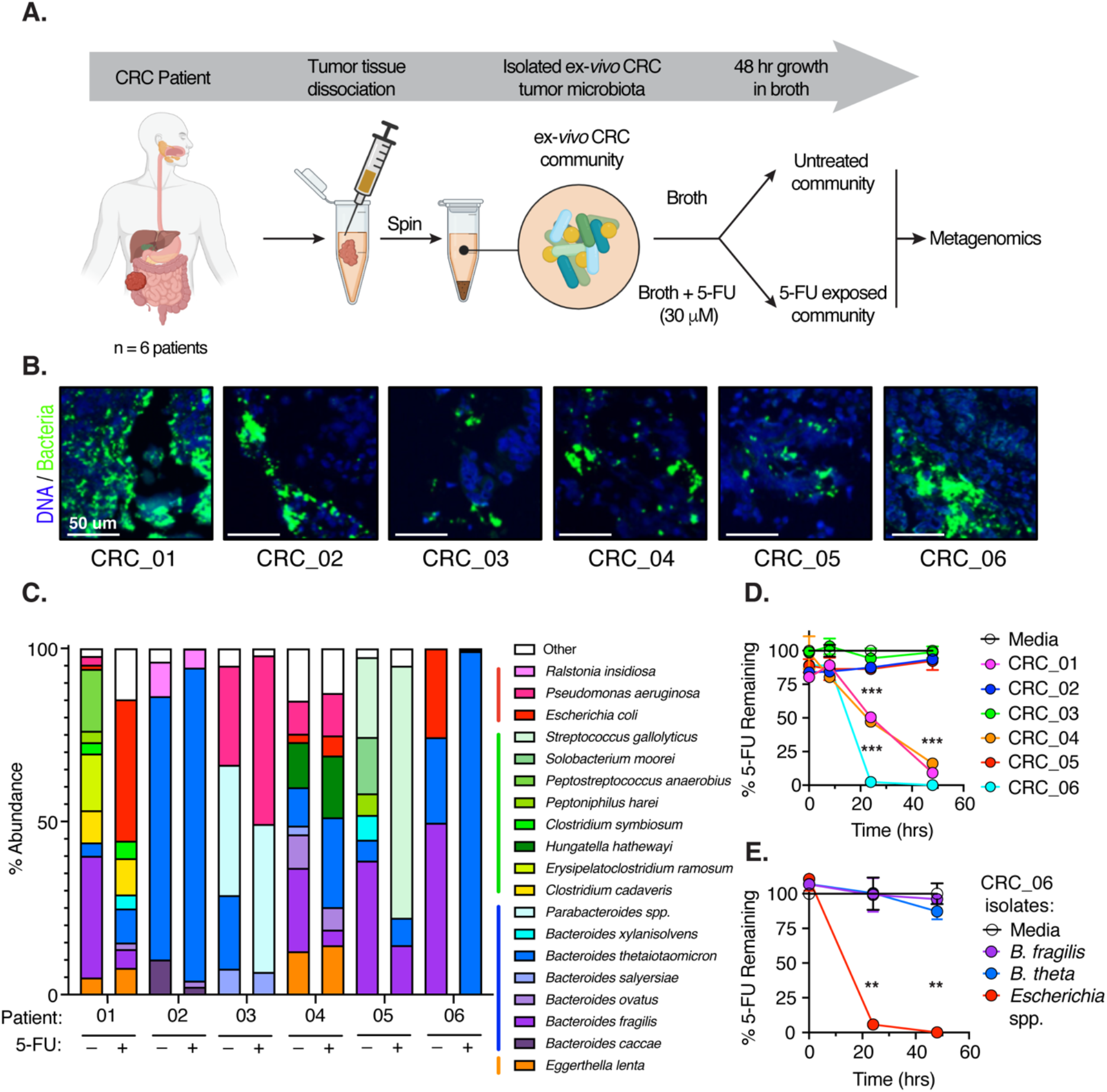
Patient-derived ex-*vivo* CRC microbiota can deplete 5-fluorouracil and reduce chemotherapeutic toxicity. (A) Schematic depicting the workflow of patient tissue processing for bacterial community isolation and downstream 5-FU exposure for 48 hours followed by metagenomic sequencing. (B) RNAscope-based fluorescence in situ hybridization (FISH) of CRC patient tumor tissue (n=6 patients). Color key: eubacterial 16S rRNA (green) and DNA (blue). (C) Relative abundance of bacterial species greater than 1% in their respective tissue samples (n=6 patients). The exposure to 5-FU (30 μM) is indicated for each community. (D) Measurement of 5-FU (4μM) disappearance in the supernatant when exposed to the indicated bacterial strains or media alone for 0, 8, 24, and 48 hours. 100% is set to the media-alone condition at 0 hrs. (E) Measurement of 5-FU (4μM) disappearance in the supernatant when exposed to the indicated bacterial strains isolated from patient CRC_06 or media alone for 0, 24, and 48 hours. 100% is set to the media-alone condition at 0 hrs. For D, E: ** indicates p-values <0.01, and ** indicates p-values <0.001 as determined by a Student’s t-test.

However, some patient community structures appeared relatively stable, which suggests resilience of these populations toward 5-FU toxicity (patients CRC_02, CRC_03, CRC_04) (Figure 4C and Table S3). The bacterial species that increased in % abundance after 5-FU exposure included *E. coli, Bacteroides thetaiotaomicron, Pseudomonas aeruginosa*, and *Streptococcus gallolyticus*. To determine if any of these ex-*vivo* CRC communities can impact 5-FU availability, we assessed 5-FU levels in the presence of these patient’s *ex-vivo* microbiota over 48 hours. We observed rapid 5-FU depletion in 50% of patient *ex-vivo* CRC communities (Figure 4D), with the tumor microbiota from patient CRC_06 (Figure 4D) being the most efficient at removal of 5-FU from media. To delineate the species responsible for 5-FU modification in patient CRC_06, we cultured isolates of the three most abundant species within the community (*Bacteroides fragilis, B. thetaiotaomicron*, and an *Escherichia* spp.) and again assessed 5-FU levels over time via LC-MS. In agreement with our previous observation, an isolate from the genus *Escherichia* was responsible for 5-FU disappearance (Figure 4E). These results offer support to the hypothesis that in a subset of CRC patients the intratumoral microbiota harbors the ability to deplete 5-FU, which could lower chemotherapeutic efficacy within colonized microniches. Importantly, 5-FU reduction would protect both sensitive oncogenic bacteria such as *Fn* and resident cancer cells from the drugs toxicity.

## Discussion

Bacterial members of the intratumoral microbiota are metabolically active in CRC and, along with malignant cells, interact with the anti-metabolite chemotherapeutic 5-FU. Our analyses reveal that interactions between the CRC tumor microbiota and 5-FU are highly complex and bacterial community members appear to fit into three distinct categories in relation to the drug: highly sensitive (e.g. *Fn*), resistant (e.g. *B. fragilis, B. breve*, and *P. micra*), and resistant and depleting (e.g. *E. coli*). Analysis of community composition in *ex-vivo* CRC microbiota after exposure to 5-FU demonstrates there is considerable loss of species diversity in a subset of CRC patient communities; suggesting that in addition to *Fn*, other bacterial species may be sensitive to 5-FU. This is supported by a recent preprint assessing 5-FU sensitivity and resistance within a range of bacterial species, albeit not CRC isolates^32^. Conversely, bacterial species present in CRC tumors have the potential to internalize and detoxify 5-FU, likely through dedicated nucleoside import and pyrimidine scavenging pathways^33–37^.

*Fn* is highly enriched in CRC tissue and multiple studies indicate that *Fn* is pathologically and clinically associated with cancer recurrence and patient outcomes^14,16,18,38^.

We found that 5-FU has potent antibacterial activity against CRC tumor isolates of *Fn* (n=11), with an average IC50 in the nanomolar range (720 nM), indicating that 5-FU-based chemotherapeutic treatment could inhibit the growth of this oncomicrobe within patient tumors. As a chemotherapeutic, 5-FU provides the greatest efficacy in CRC compared to other cancer types^19^, raising the intriguing possibility that 5-FU treatment efficacy could in part be due to its unanticipated role as a potent antimicrobial agent against dominant members of the intratumoral microbiota, including *Fn*.

The presence of *Fn* is also associated with cancer recurrence after treatment. A study by Yu *et al*.^14^ found that *Fusobacterium* was consistently enriched and had higher bacterial loads in recurrent CRC tissues compared to non-recurrent CRC tissues across multiple patient cohorts. In a congruent study, Serena *et al.^16^* detected *Fn* in 58% of treatment-naïve tumor biopsies from patients (n=143) with locally advanced rectal cancer and discovered *Fn* persists in 26% of tumors treated with 5-FU based neoadjuvant chemo-radiotherapy (nCRT)^16^. *Fn*-positivity in tissue after nCRT, but not its baseline status in tissue, was significantly correlated with a 7.5 fold increased risk of relapse in patients^16^. This work is in agreement with both studies; we expect that the subset of patients where *Fn* survives 5-FU toxicity likely contain an intratumoral bacterial community capable of metabolizing 5-FU, protecting both cancer and sensitive bacterial cells such as *Fn*, promoting CRC recurrence. To expand upon this, prospective studies that investigate tumor microbiomes of biopsy’s pre-chemotherapy, and resections post-chemotherapy, in addition to patient treatment response are warranted.

We identify multiple strains of *E. coli* that lead to 95% depletion of available 5-FU as early as 24 hours post-exposure (Figure 3B, Figure 4E, and Figure S3B). *E. coli*-exposed 5-FU no longer induces CRC epithelial cell growth inhibition, and this treatment is protective of *Fn* isolated from tumors. These results complement research from two recent studies that reported 5-FU efficacy was altered in *Caenorhabditis elegans* that were fed genetically different bacterial strains of *E. coli*^39,40^. In humans, 5-FU is detoxified through metabolic conversion to dihydrofluorouracil (DHFU) by the protein dihydropyrimidine dehydrogenase (DPD)^41,42^. Prior research has demonstrated that *E. coli* harbors a functional homolog of human DPD within an operon called *pre*TA that can convert uracil into 5,6-dihydrouracil in *vivo*^43^. Additionally, a recent preprint by Spanogiannopoulos *et al*.^32^, demonstrates that bacterial homologs of PreTA, including those in *E. coli*, can metabolize 5-FU to DHFU *in vitro*. Since *E. coli* is prevalent in CRC patient tumor tissue^29^, it has the potential to contribute to the local depletion of 5-FU and repression of its toxicity on otherwise-sensitive cells in the tumor. This work is consistent with a growing body of literature supporting that microbiota composition has a dramatic impact (both directly and indirectly) on the chemotherapeutic efficacy of many drugs in a wide variety of cancers ^1,44^ It was recently determined that many members of the Gammaproteobacteria class directly metabolize and detoxify gemcitabine, a primary chemotherapeutic for the treatment of pancreatic ductal adenocarcinoma, leading to gemcitabine resistance in murine models^45^. Collectively, these findings support that the composition of microbiota might inform the design of patient treatment regimens in CRC.

Patients treated with 5-FU based chemotherapy for CRC undergo multiple rounds of treatment, with intermittent breaks of two to four weeks to allow for recovery. Currently, there is no standard dosing protocol beyond body surface area for 5-FU in patients ^46^ and administered levels of 5-FU are often adjusted throughout treatment. There is a subset of patients that do not respond to 5-FU or develop chemoresistance to the drug during treatment^47^. Our *in vitro* work suggests that microbiota-associated modification of 5-FU might contribute to patient 5-FU chemoresistance. In humans, changes in expression of thymidylate synthase, nucleoside importers, and multidrug efflux pumps contribute to chemoresistance to 5-FU in patients^47^. Given that these classes of proteins also exist in many bacterial species^28,34,48,49^, repeat exposure selects for mutations in these pathways leading to 5-FU tolerance^32,50^. These findings support the benefits of taking the tumor-associated microbiota into consideration when stratifying patients into risk-categories for 5-FU resistance especially in the setting of neoadjuvant chemotherapy prior to tumor resection.

## Methods

### Bacterial culture

All bacteria were grown from cryostocks on fastidious anaerobe agar plates (FAA; Neogen) supplemented with 10% defibrinated horse blood (DHB; Lampire Biological Laboratories). For *Fusobacterium* culture FAA+10% DHB plates included josamycin (3 μg/mL), vancomycin (4 μg/mL), and norfloxacin (1 μg/mL) for selective culturing (JVN). For liquid-based assays, bacteria were incubated in either tryptic soy broth (TSB; Sigma Aldrich), brain heart infusion broth (BHI; Research Products International), or fastidious anaerobe broth (FAB; Neogen). All strains were stored at −80°C in TSB supplemented with 40% (v/v) glycerol. All bacterial culturing occurred under anerobic conditions in an anaerobic chamber (Anaerobe Systems AS-580) or sealed box with an AnaeroGen gas pack (Oxoid) and incubated at 37°C. *Fn* isolates, *P. micra, B. fragilis*, and *B. breve* were cultured for 2 days on FAA + 10% DHB prior to assay inoculation. *E. coli* was cultured for 1 day on FAA + 10% DHB prior to assay inoculation. List of strains used in this study can be found in **Table S4**.

### Mammalian cell culture

Human colon cancer epithelial RKO cells (ATCC CRL-2577) were grown in McCoys 5A with L-glutamine (Corning) supplemented with 10% (v/v) fetal bovine serum (Sigma). Antibiotics (100 U/mL penicillin, 100 μg/mL streptomycin; Gibco) were included in the growth medium during maintenance of cell lines and removed 24 hours prior to bacterial co-culture. Cell cultures were incubated at 37°C in 5% CO_2_

### Bioactive library screen and dose-response validation

#### Bioactive library screen

1,846 compounds were printed in duplicate on the bottom of 384 well plates by the Broad Therapeutics Platform (**Table S1**). The addition of 30 μl of liquid in each well results in 32 μM compound. Each plate had multiple neutral controls (TSB only) and inhibitory controls (5 μg/mL metronidazole in TSB).

#### Dose-response validation of inhibitory compounds and antineoplastic agents

384 well plates printed with eight concentrations of each compound in duplicate (30, 15, 7,5, 3.75, 1.88, 0.94, 0.47, and 0.23 μM). These plates included the 34 inhibitory compounds confirmed in both biological replicates of our pilot screen and an additional 21 antineoplastic agents of interest (**Table S2**). Each plate had multiple neutral controls (TSB only) and inhibitory controls (5 μg/mL metronidazole in TSB).

#### Plate inoculation and viability determination

Each well was inoculated with 30 μl of 2×10^6^ colony forming units (CFU) per mL of *Fna* SB010 resuspending in TSB. Plates were incubated anaerobically at 37°C for 48 hrs. Cells were lysed via the addition of 10 μl BacTiter Glo (Promega) and ATP levels measured through luminescence detection on a PerkinElmer Envision. Spectrometry parameters: read time=0.1 sec, plate height= 0 cm.

#### Data analysis

The inhibitor hit cut-off was set at three standard deviations from the mean of the neutral control (−86%; broth + DSMO). The data was normalized to the DMSO only and DMSO - inhibitor control (metronidazole). The data generated in the dose-response curves was normalized to the DMSO – inhibitor control.

### Bacterial dose-response curves

5-fluorouracil (5-FU; >99% purity; Tokyo Chemical Industry) stocks were stored as 5 mg/mL aliquots in dimethyl sulfoxide (DMSO) at 4°C for up to one month. Dose-response plates were set up through the dilution of 5 mg/mL 5-FU stock in BHI +0.1% DMSO to the desired upper concentration, then serially diluted 2-fold in BHI + 0.1% DMSO until the desired lower concentration was reached. The lowest 5-FU concentration used was 0.0375 μM and the highest concentrations reached 4.8 to 38.4 μM 5-FU for 8 pt and 11 pt dose response curves respectively. The control for positive inhibition (0% expected growth) for *Fn* isolates, *B. fragilis*, and *B. breve* was 5 μg/mL metronidazole + 0.1% DMSO in BHI. The positive inhibition control for *P. micra* and *E. coli* was 30 μg/mL gentamycin in BHI + 0.1% DMSO. The negative inhibition control (100% growth) was the base BHI + 0.1% DMSO media. Every plate tested included additional no bacteria controls wells of 5-FU, metronidazole, gentamycin, and broth alone to monitor for any sources of external bacterial contamination. Each well (100 μl volume) was inoculated with 5 μl of 1×10^8^ CFU/mL of *Fn and P. micra* (5×10^6^ CFU/mL; 5×10^4^ CFU/well) or 5 μl of 2×10^7^CFU/mL *B. fragilis, B. breve*, and E. coli (1×10^6^ CFU/mL; 1×10^4^ CFU/well). Plates were incubated in a 2.5L sealed box at 37°C for 48 hours (hrs) under anaerobic conditions using an AnaeroGen gas pack. Bacterial viability was determined through ATP level measurements using a BacTiter Glo microbial cell viability assay (Promega). Cells were lysed through addition of 33 μl BacTiter Glo and ATP levels were measured through luminesce quantification on a spectrophotometer (BioTek Synergy H4) using the following parameters: Gain: 135, Integration Time: 1 sec, Read Height: 4.5 mm).

### Bacterial co-culture with 5-FU and the generation of “modified” 5-FU supernatants

#### For Fn protection or mass-spectrometry assays

4 μM 5-FU was incubated with 5×10^8^ CFU/mL bacteria in a total volume of 1 mL of broth (TSB for *Fn* protection assays and BHI for mass spectrometry) for 48 hrs at 37°C. For patient-derived ex-*vivo* CRC communities, initial communities were diluted 1/50 in 4 μM 5-FU in 1 mL of FAB for 48 hrs. Broth media + 400 μM 5-FU (100x) was incubated for 48 hrs at 37°C in order to generate the “fresh” 5-FU control.

#### For use in mammalian cell culture assays

20 μM 5-FU was incubated with 5×10^8^ CFU/mL bacteria in a total volume of 1 mL McCoys 5A + L-glutamine+ 10% FBS for 48 hrs. Cells were then pelleted at 7000 × g for 3 min and resulting supernatants filtered through a 0.2 μm filter. Media alone with 2 mM 5-FU (100×) was incubated for 48 hrs at 37°C in order to generate the “fresh” 5-FU control.

#### Supernatant generation

Cells were then pelleted at 7000 × g for 3 min and resulting supernatants filtered through a 0.2 μm PVDF filter (Millipore). The no bacteria “fresh” 5-FU controls were also filtered in this way to account for any potential loss of 5-FU by filtration. For the “fresh” 5-FU conditions, bacterial supernatants (500 μl) that had grown in broth alone were spiked with the 100X “fresh” 5-FU control (5 μl).

### *Fn* “modified” 5-FU protections assays

#### Supernatant protection

5×10^6^ CFU/mL *Fna* SB010 was grown in *B. fragilis, E. coli* CRC, or *E. coli pks+* supernatants incubated in TSB alone, TSB + 4 μM 5-FU (modified), or TSB alone supplemented with 4 μM 5-FU after bacterial filtration (fresh). *Fn* was grown for 48 hrs at 37°C under anaerobic conditions, and bacterial viability was measured through ATP determination with BacTiter Glo described in detail in the “bacterial dose-response curves” methods section. 100% growth was set to *Fna* SB010 viability in TSB alone, or in each bacterial supernatant along without the addition of 5-FU.

#### Direct co-cultureprotection

5×10^6^ CFU/mL *Fna* SB010 was incubated with 4 μM 5-FU alone or with the addition of 1×10^6^ CFU/mL *E. coli* CRC for 0, 24, and 48 hrs at 37°C under anaerobic conditions. As a control, *Fna* SB010 was also incubated in media alone or with 1×10^6^ CFU/mL *E. coli* CRC alone without the addition of 5-FU. Bacterial viability was monitored at 0, 24, and 48 hours by 10-fold serial dilutions and plating for CFU. *Fna* SB010 colonies were isolated through plating on FAA + 10% DHB + 30 μg/mL gentamycin and incubating under anaerobic conditions. *E. coli* CRC colonies were isolated through plating on FAA + 10% DHB and incubating under aerobic conditions.

### CRC epithelial cell “modified” 5-FU protection assays

#### Supernatant protection

RKO cells were seeded into a clear bottom white-96 well plate at a density of 5000 cells/well (100 μl total volume) and incubated for 16 hrs at 37°C to allow for surface attachment. Filtered bacterial supernatants 48 hours incubation with media alone (McCoys 5A), 20 μM 5-FU (modified), or media alone supplemented with 20 μM 5-FU after bacterial filtration (fresh) were added to RKO cells cell culture media in a 1:4 dilution (33.33 μl) for an anticipated final concentration of 5 μM 5-FU (the reported IC50 for RKO cells). RKO cells were then incubated for 72 hrs at 37°C in an IncuCyte S3 live imaging incubator (Sartorius and Essun BioScience). Each well was imaged in five locations with a 10X objective every two hrs for a total of 72 hrs (Camera: Basler Ace 1920-155μm CMOS). RKO cell growth over time was determined by calculating the % confluence of each well after training the IncuCyte Live Cell-Analysis software on RKO cell parameters and filtering out debris (volumes <150 μm^3^).

### RNAscope-based fluorescent in situ hybridization

Paraffin embedded sections (5μm) were deparaffinized, pretreated, and hybridized using RNAscope Multiplex Fluorescent Assay V2 (Advanced Cell Diagnostics, Inc., Newark, CA, USA) in accordance with the manufacturer’s protocol. The probe used was EB-16s-rRNA targeting a conserved region of 16s rRNA of all bacteria. Probes were hybridized for two hours then washed twice in 1X RNAscope Wash Buffer Solution for 2 minutes. Three amplification steps were performed, then probe channels we developed and stained individually. The Eubacterium probe was stained with Opal 690 and samples were counterstained with DAPI for 30 seconds, prior to mounting in ProLong Gold (Thermofisher Scientific) Whole slide scanning was performed on a Vectra Polaris multispectral imaging system (Akoya Biosciences Inc.) at 40× (25 μm/pixel). Sample and corresponding negative control probe slides were scanned under the same exposure conditions.

### 5-FU analysis through mass spectrometry

5-FU from bacterially modified media samples was analyzed by liquid-chromatography with tandem quadrupole mass spectrometry (LC-MS/MS QQQ) using a Waters Xevo-XS with a Waters Acquity I-Class UPLC. Samples were mixed 1:1 with 0.4 μM of a heavy, stable isotope of 5-FU in diH_2_O (98 atom %^15^N, 99 atom %^13^C; empirical forumla: ^13^CC_3_H_3_F^15^N_2_O_2_; Millipore Sigma) that acted as an internal standard for 5-FU quantitation. Chromatographic separation was achieved using a Thermo Hypercarb column (2.1×50 mm, 3u) at room temperature. The mobile phase consisted of solution A: 100 mM NH_4_OAC in H_2_O, pH 9.5 using NH4OH, and solution B: 0.1% NH_4_OH in acetonitrile. A flow rate of 0.3 mL/min was used with the following gradient elution profile: 100% A from 0-6 minutes, 50% A 50% B from 6-7.1 minutes, 100% A from 7.1-10 minutes. 50 μl of sample was injected for LC-MS/MS QQQ analysis. This assay used multiple reaction monitoring (MRM) in electrospray negative ionization, and the ions measured for the quantitation were: 5-Flourouracil: 129.03->42.16 & 129.03->129.0 and C13-5-Flourouracil: 132.1->60.1 & 132.1->44.1. Quantitative analysis for 5-FU was performed using an isotope dilution method with a 7 pt calibration curve (4-fold dilutions starting at 20 μM) run in duplicate with a linear regression coefficient of R^2^= 0.9998. The level of quantitation achieved was 0.078 μM and spanned to 20 μM.

### Ex-*vivo* CRC tumor microbiota generation

#### Patient cohort

The six patient specimens included in this study were collected under Fred Hutchinson Cancer Research Center IRB protocol RG1006974. All six patients were treatment naive and tissue was collected during tumor resection.

#### Tissue dissociation

Frozen patient tumor tissue was thawed at 37°C until only amount of ice was left. Tissue was transferred to a sterile petri dish and minced with scalpels. Tissue was then transferred to a 5 mL conical tube and covered with 2.5 mL of FAB and vortexed for 30 seconds. Tissue was then manually ground against the bottom of the conical tube with a 5 mL serological pipette. Tissue was then passed through a 16-gauge needle until there was a uniform suspension, followed by an 18-gauge needle. Cells were centrifuged at 300xg for 4 minutes, and the top 2 mL of supernatant was moved into a 2 mL eppitube. The eukaryotic cell pellet and remaining 500 μl was cryopreserved for downstream microbial culturing. From this 2 mL supernatant that contained the ex-*vivo* tissue microbiota, 1 mL as removed and centrifuged at 7,000×g for 3 minutes, and the pellet stored at −20°C as marker of the initial microbial population prior to growth in broth. The remaining 1 mL of supernatant was divided into two tubes (500 ul each) and supplemented to 500 μl of FAB. One tube was supplemented with 50 μl of 600 μM 5-FU (final 30 μM). Both tubes were transported to an anaerobic chamber, opened to allow gas exchange, the sealed and incubated for 48 hours at 37°C. After 48 hours, 100 μl was removed, diluted 1:5 in cryoprotectant and stored at −80°C for downstream microbial culturing (eg. 5-FU disappearance assays). The remaining 900 μl was centrifuged at 7,000×g for 3 minutes, the supernatant removed, and the pellets stored at −20°C for future 16S rRNA sequencing and metagenomic analysis.

### Shotgun metagenomic sequencing

The samples were processed and analyzed with the ZymoBIOMICS®Shotgun Metagenomic Sequencing Service for Microbiome Analysis (Zymo Research, Irvine, CA).

#### DNA Extraction

One of three DNA extraction kits was used depending on the sample type and sample volume. In most cases, the ZymoBIOMICS®-96 MagBead DNA Kit (Zymo Research, Irvine, CA) was used to extract DNA using an automated platform. In some cases, ZymoBIOMICS®DNA Miniprep Kit (Zymo Research, Irvine, CA) was used. For some low biomass samples, such as skin swabs, the ZymoBIOMICS®DNA Microprep Kit (Zymo Research, Irvine, CA) was used as it permits for a lower elution volume, resulting in more concentrated DNA samples.

#### Shotgun Metagenomic Library Preparation

Genomic DNA samples were profiled with shotgun metagenomic sequencing. Sequencing libraries were prepared with either the KAPA™HyperPlus Library Preparation Kit (Kapa Biosystems, Wilmington, MA) with up to 100 ng DNA input following the manufacturer’s protocol using internal single-index 8 bp barcodes with TruSeq®adapters (Illumina, San Diego, CA) or the Nextera®DNA Flex Library Prep Kit (Illumina, San Diego, CA) with up to 100 ng DNA input following the manufacturers protocol using internal dual-index 8 bp barcodes with Nextera^®^ adapters (Illumina, San Diego, CA). All libraries were quantified with TapeStation®(Agilent Technologies, Santa Clara, CA) and then pooled in equal abundance. The final pool was quantified using qPCR.

#### Sequencing

The final library was sequenced on either the Illumina HiSeq®or the Illumina NovaSeq®.

#### Bioinformatics Analysis

Raw sequence reads were trimmed to remove low quality fractions and adapters with Trimmomatic-0.33 (Bolger et al., 2014): quality trimming by sliding window with 6 bp window size and a quality cutoff of 20, and reads with size lower than 70 bp were removed. Antimicrobial resistance and virulence factor gene identification was performed with the DIAMOND sequence aligner (Buchfink et al., 2015). Microbial composition was profiled with Centrifuge (Kim et al., 2016) using bacterial, viral, fungal, mouse, and human genome datasets. Strain-level abundance information was extracted from the Centrifuge outputs and further analyzed: (1) to perform alpha- and beta-diversity analyses; (2) to create microbial composition barplots with QIIME (Caporaso et al., 2012); (3) to create taxa abundance heatmaps with hierarchical clustering (based on Bray-Curtis dissimilarity); and (4) for biomarker discovery with LEfSe (Segata et al., 2011) with default settings (p>0.05 and LDA effect size >2).

### Statistics

All statistical tests were performed in GraphPad Prism 7.0 with α=0.05; p<0.05 is indicated by asterisks with the following key: *<0.05, **<0.01, ***<0.001

## Acknowledgements

We thank Brian Reid for providing primary CRC patient tissue, Jim Berstler and Kate Hartland from the Broad Center for the Development of Therapeutics (CDoT) for support and analysis of the small molecule screens, Dale Whittington from the University of Washington Medicinal Chemistry Mass Spectrometry Center for assistance in 5-FU detection and depletion protocol development, and the Histology Core at the Fred Hutchinson Cancer Research Center for tissue sectioning and fluorescent scanning.

## Author Contributions

K.D.L., S.B and C.D.J designed the study. K.D.L, A.B., A.K., and S.B. performed the experiments, and K.D.L and S.B. analyzed data. K.D.L., C.D.J., and S.B. wrote the manuscript.

---

**Figure S1:**
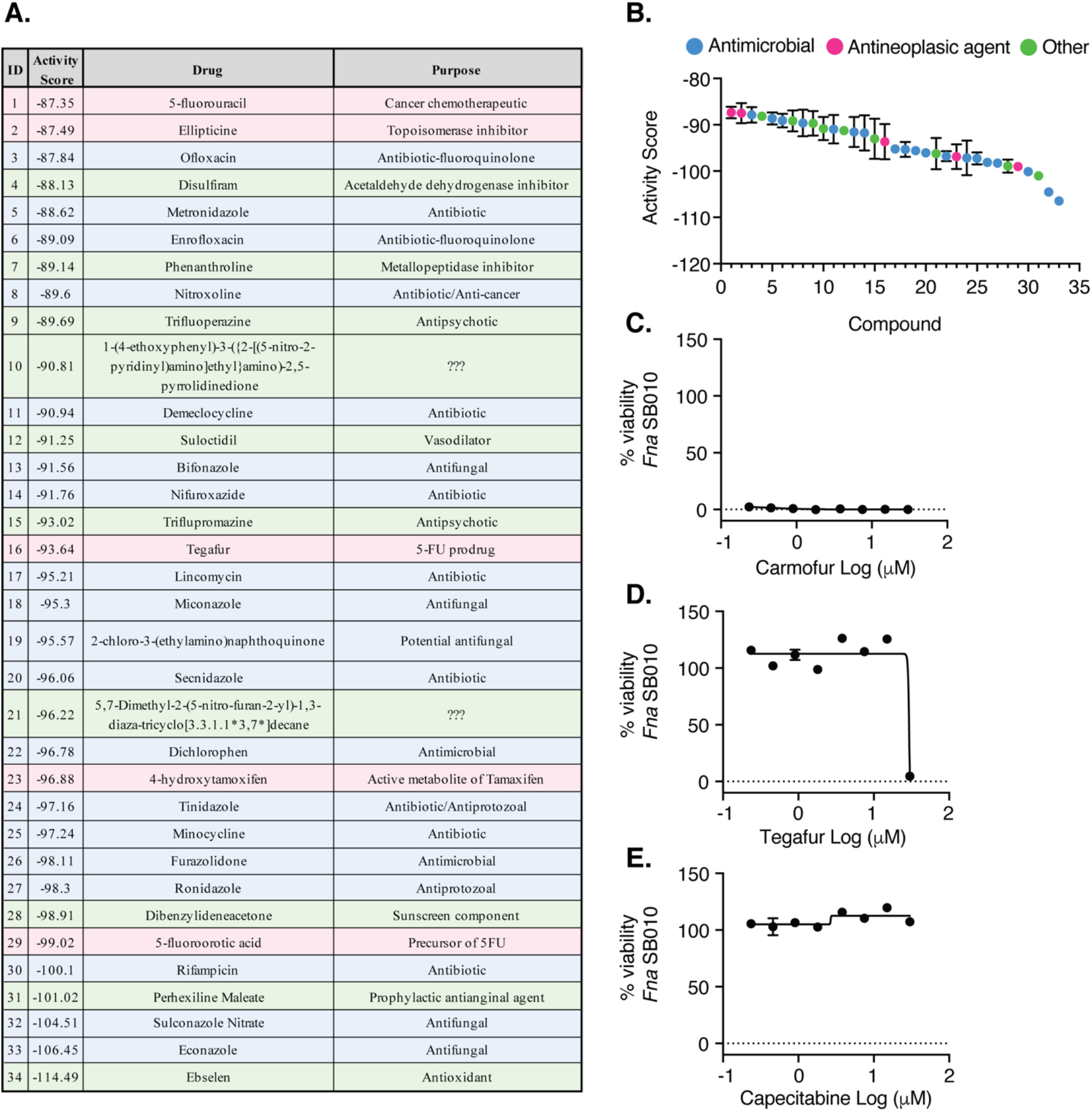
List of 34 *F. nucleatum* growth inhibitors and dose-response to prodrugs of 5-fluorouracil. (A) List of *F. nucleatum* growth inhibitors (n=38), their respective activity scores and function (when known). (B) Activity scores for n=34 *F. nucleatum* inhibitors. Color key for drug categories: antimicrobial (blue), antineoplastic (pink), other (green). (C-E) Eight-point dose-response curves of *F. nucleatum* viability after 48 hours of exposure to 0.25-32 μM of the indicated prodrugs of various chemotherapeutics used to treat metastatic colon cancer. The connecting line is a non-linear regression of the log(inhibitor) vs. response with a variable slope (four-parameters).

**Figure S2:**
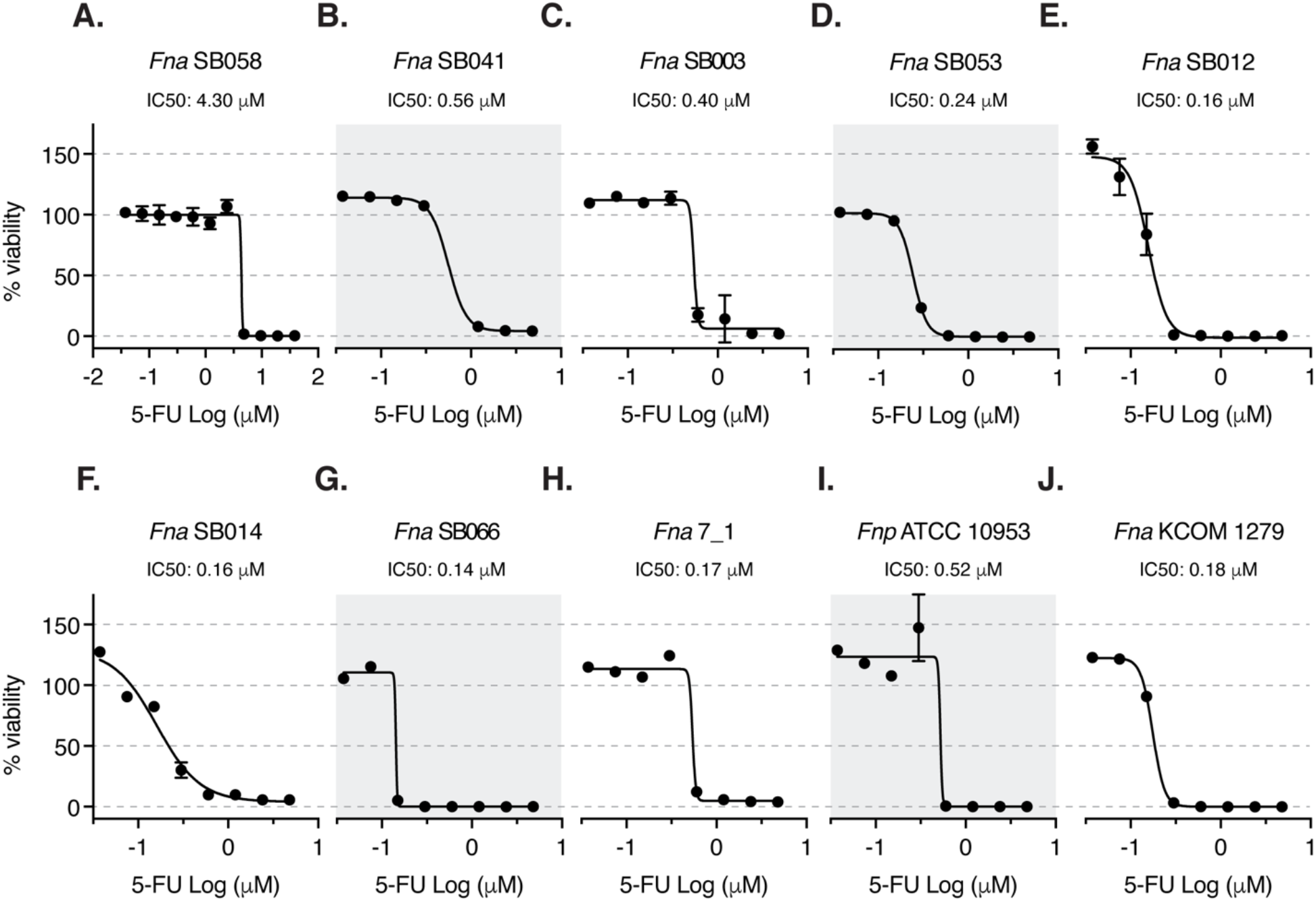
5-FU sensitivity of *F. nucleatum* isolates to 5-fluorouracil. (A) Eleven-point and (B-J) Eight-point dose response curves of relative viability of indicated *F. nucleatum* isolates after exposure to either 0.0375-38.4 μM (A) or 0.0375-4.8 μM (B-J) 5-FU for 48 hrs. The connecting line is a non-linear regression of the log(inhibitor) vs. response with a variable slope (four-parameters). The IC50 calculations are indicated above each graph.

**Figure S3:**
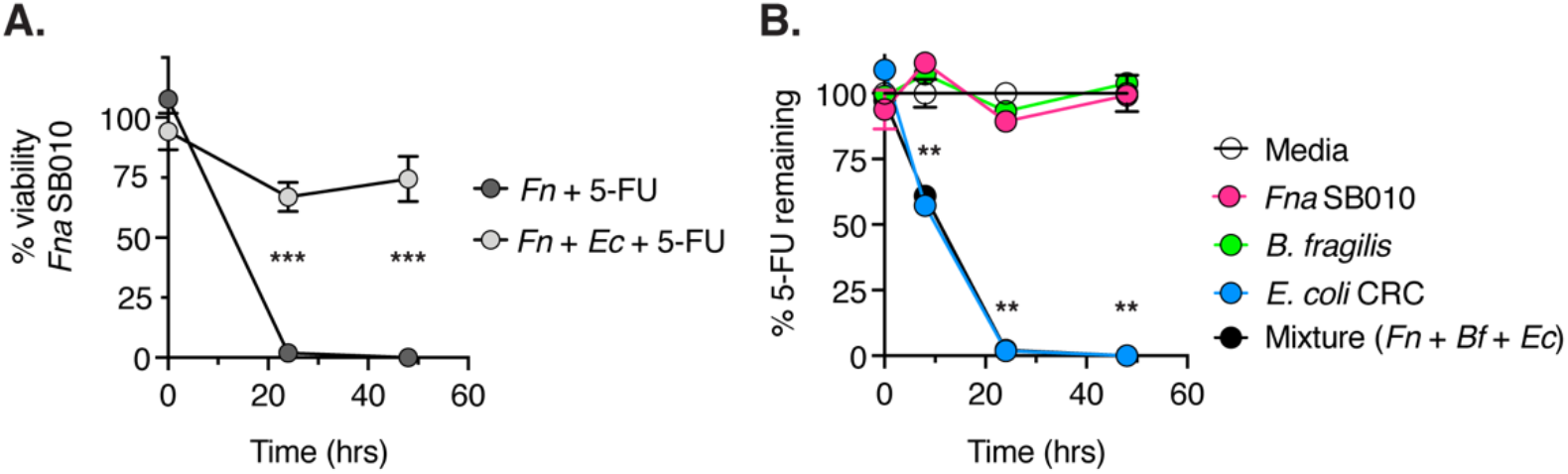
*E. coli* modifies 5-fluorouracil in co-culture with *F. nucleatum*. (A) Relative viability of *Fna* SB010 in co-culture with *E. coli* CRC (light grey circles) and 4 μM 5-FU or *Fna* SB010 alone (dark grey circles) incubated with 4 μM 5-FU alone at 0, 24, and 48 hours. Viability was determined through colony forming unit (CFU) analysis. 100% viability was set to the C.F.U. of *Fna* SB010 in either media alone or in co-culture with *E. coli* CRC str 2 alone without the addition of 5-FU at each timepoint. (B) Measurement of 5-FU (4μM) disappearance in the supernatant when exposed to *Fna* SB010, *B. fragilis*, or *E. coli* CRC to the indicated bacterial strains isolated from patient CRC_06 or media alone for 0, 8, 24, and 48 hours. 100% is set to the media-alone condition at 0 hrs. For A,B: ** indicates p-values <0.01, and ** indicates p-values <0.001 as determined by a Student’s t-test in comparison to the media only condition at each respective timepoint.

**Figure S4:**
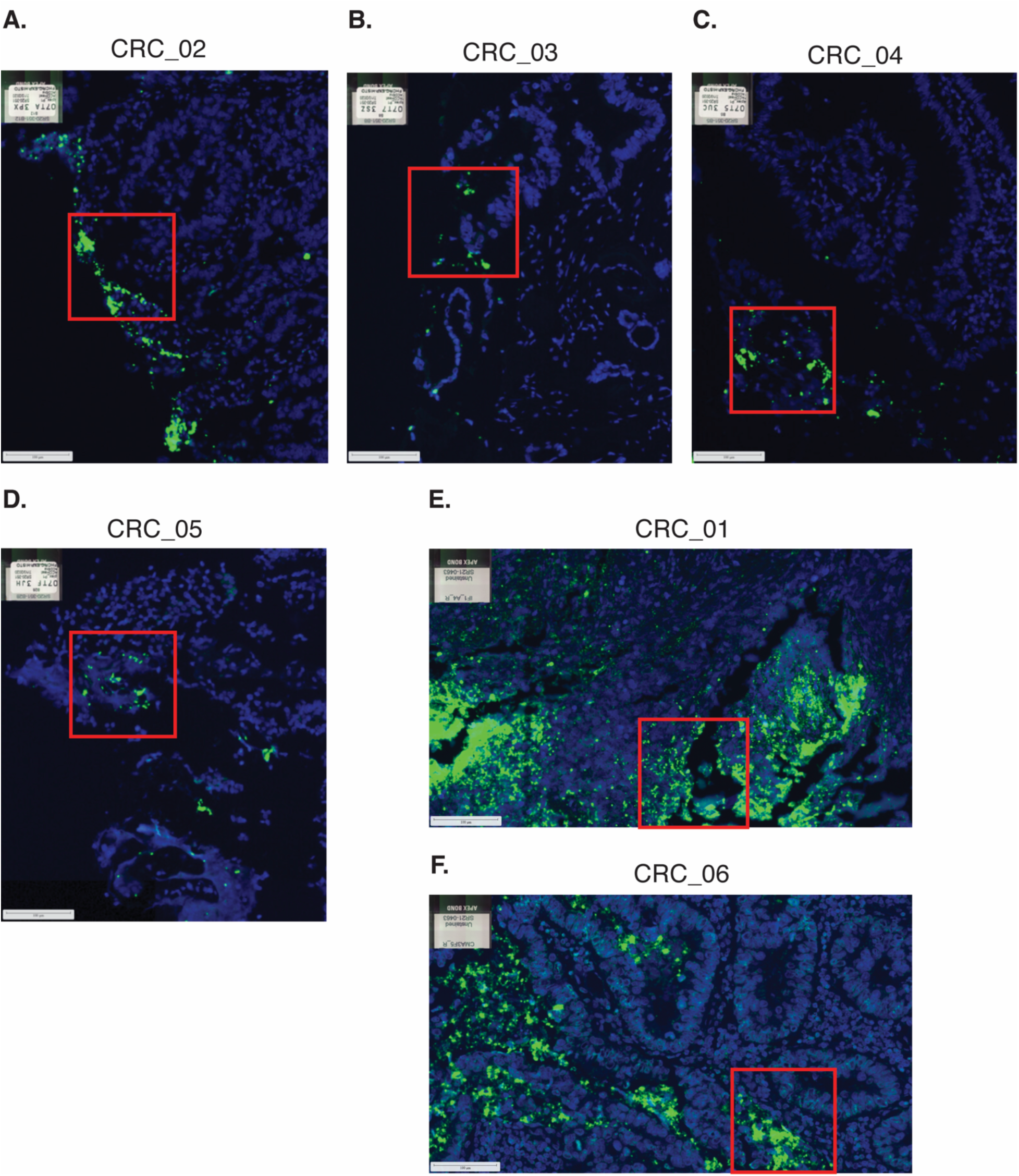
Original RNA-scope FISH imaging for patient tissues used in this study. (A-F) RNAscope-based fluorescence in situ hybridization (FISH) of CRC patient tumor tissue (n=6 patients). The red boxes indicate which region of interest was depicted in Figure 4. Color key: eubacterial 16S rRNA (green) and DNA (blue).

**Table S4.** Strains used in this study.

| Organism | Subspecies and strain ID | Reference |
| --- | --- | --- |
| *F. nucleatum*<br>subsp. *animalis* | SB003 | This study |
|  | SB010 | Bullman *et al.*, 2017<sup>18</sup> |
|  | SB012 | This study |
|  | SB014 | This study |
|  | SB041 | This study |
|  | SB053 | This study |
|  | SB058 | This study |
|  | SB066 | This study |
|  | 7_1 | Strauss *et al.*, 2011<sup>51</sup> |
|  | KCOM1279 | Korean Collection for Oral Microbiology |
| *F. nucleatum*<br>subsp. *nucleatum* | SB011 | This study |
| *F. nucleatum*<br>subsp. *polymorphum* | SB013 | This study |
|  | ATCC 10953 | ATCC |
| *F. nucleatum*<br>subsp. *vincentii* | SB054 | This study |
| *E. coli* | CRC | This study |
|  | *pks+* ATCC 25922 | ATCC |
|  | CRC_06 | This study |
| *B. fragilis* | CTX25T | This study |
|  | CRC_06 | This study |
| *B. thetaiotaomicron* | CRC_06 | This study |
| *B. breve* | Reid K10D6 | This study |
| *P. micra* | CRC_02 | This study |

## References

1 LaCourse KD, Johnston CD, Bullman S. The relationship between gastrointestinal cancers and the microbiota. Lancet Gastroenterol Hepatol 2021; 6: 498–509.

2 Kostic AD, Gevers D, Pedamallu CS et al. Genomic analysis identifies association of Fusobacterium with colorectal carcinoma. Genome Res 2012; 22: 292–298.

3 Castellarin M, Warren RL, Freeman JD et al. Fusobacterium nucleatum infection is prevalent in human colorectal carcinoma. Genome Res 2012; 22: 299–306.

4 McCoy AN, Araújo-Pérez F, Azcárate-Peril A, Yeh JJ, Sandler RS, Keku TO. Fusobacterium is associated with colorectal adenomas. PloS One 2013; 8: e53653.

5 Tahara T, Yamamoto E, Suzuki H et al. Fusobacterium in Colonic Flora and Molecular Features of Colorectal Carcinoma. Cancer Res 2014; 74: 1311–1318.

6 Flanagan L, Schmid J, Ebert M et al. Fusobacterium nucleatum associates with stages of colorectal neoplasia development, colorectal cancer and disease outcome. Eur J Clin Microbiol Infect Dis Off Publ Eur Soc Clin Microbiol 2014; 33: 1381–1390.

7 Ito M, Kanno S, Nosho K et al. Association of Fusobacterium nucleatum with clinical and molecular features in colorectal serrated pathway. Int J Cancer 2015; 137: 1258–1268.

8 Li Y-Y, Ge Q-X, Cao J et al. Association of Fusobacterium nucleatum infection with colorectal cancer in Chinese patients. World J Gastroenterol 2016; 22: 3227–3233.

9 Kostic AD, Chun E, Robertson L et al. Fusobacterium nucleatum Potentiates Intestinal Tumorigenesis and Modulates the Tumor-Immune Microenvironment. Cell Host Microbe 2013; 14: 207–215.

10 Rubinstein MR, Wang X, Liu W, Hao Y, Cai G, Han YW. Fusobacterium nucleatum Promotes Colorectal Carcinogenesis by Modulating E-Cadherin/β-Catenin Signaling via its FadA Adhesin. Cell Host Microbe 2013; 14: 195–206.

11 Yang Y, Weng W, Peng J et al. Fusobacterium nucleatum Increases Proliferation of Colorectal Cancer Cells and Tumor Development in Mice by Activating TLR4 Signaling to NFκB, Upregulating Expression of microRNA-21. Gastroenterology 2017; 152: 851–866.e24.

12 Yu Y-N, Yu T-C, Zhao H-J et al. Berberine may rescue Fusobacterium nucleatum-induced colorectal tumorigenesis by modulating the tumor microenvironment. Oncotarget 2015; 6: 32013–32026.

13 Chen Y, Peng Y, Yu J et al. Invasive Fusobacterium nucleatum activates beta-catenin signaling in colorectal cancer via a TLR4/P-PAK1 cascade. Oncotarget 2017; 8: 31802–31814.

14 Yu T, Guo F, Yu Y et al. Fusobacterium nucleatum Promotes Chemoresistance to Colorectal Cancer by Modulating Autophagy. Cell 2017; 170: 548–563.e16.

15 Mima K, Nishihara R, Qian ZR et al. Fusobacterium nucleatum in colorectal carcinoma tissue and patient prognosis. Gut 2016; 65: 1973–1980.

16 Serna G, Ruiz-Pace F, Hernando J et al. Fusobacterium nucleatum persistence and risk of recurrence after preoperative treatment in locally advanced rectal cancer. Ann Oncol 2020; 31: 1366–1375.

17 Zhang S, Yang Y, Weng W et al. Fusobacterium nucleatum promotes chemoresistance to 5-fluorouracil by upregulation of BIRC3 expression in colorectal cancer. J Exp Clin Cancer Res 2019; 38: 14.

18 Bullman S, Pedamallu CS, Sicinska E et al. Analysis of Fusobacterium persistence and antibiotic response in colorectal cancer. Science 2017; 358: 1443–1448.

19 Longley DB, Harkin DP, Johnston PG. 5-fluorouracil: mechanisms of action and clinical strategies. Nat Rev Cancer 2003; 3: 330–338.

20 Neugut AI, Lin A, Raab GT et al. FOLFOX and FOLFIRI use in Stage IV Colon Cancer: Analysis of SEER-Medicare Data. Clin Colorectal Cancer 2019; 18: 133–140.

21 Abe Y, Sakuyama N, Sato T et al. Evaluation of the 5-fluorouracil plasma level in patients with colorectal cancer undergoing continuous infusion chemotherapy. Mol Clin Oncol 2019; 11: 289–295.

22 Fang L, Jiang Y, Yang Y et al. Determining the optimal 5-FU therapeutic dosage in the treatment of colorectal cancer patients. Oncotarget 2016; 7: 81880–81887.

23 Matsumoto H, Okumura H, Murakami H et al. Fluctuation in Plasma 5-Fluorouracil Concentration During Continuous 5-Fluorouracil Infusion for Colorectal Cancer. Anticancer Res 2015; 35: 6193–6199.

24 Kook J-K, Park S-N, Lim YK et al. Genome-Based Reclassification of Fusobacterium nucleatum Subspecies at the Species Level. Curr Microbiol 2017; 74: 1137–1147.

25 Kook J-K, Park S-N, Lim YK et al. Fusobacterium nucleatum subsp. fusiforme Gharbia and Shah 1992 is a later synonym of Fusobacterium nucleatum subsp. vincentii Dzink et al. 1990. Curr Microbiol 2013; 66: 414–417.

26 Ye X, Wang R, Bhattacharya R et al. Fusobacterium Nucleatum Subspecies Animalis Influences Proinflammatory Cytokine Expression and Monocyte Activation in Human Colorectal Tumors. Cancer Prev Res Phila Pa 2017; 10: 398–409.

27 Borozan I, Zaidi SH, Harrison TA et al. Molecular and Pathology Features of Colorectal Tumors and Patient Outcomes Are Associated with Fusobacterium nucleatum and Its Subspecies animalis. Cancer Epidemiol Biomark Prev Publ Am Assoc Cancer Res Cosponsored Am Soc Prev Oncol 2021. doi:10.1158/1055-9965.EPI-21-0463.

28 Islam Z, Gurevic I, Strutzenberg TS, Ghosh AK, Iqbal T, Kohen A. Bacterial versus human thymidylate synthase: Kinetics and functionality. PLoS ONE 2018; 13. doi:10.1371/journal.pone.0196506.

29 Buc E, Dubois D, Sauvanet P et al. High prevalence of mucosa-associated E. coli producing cyclomodulin and genotoxin in colon cancer. PloS One 2013; 8: e56964.

30 Dejea CM, Fathi P, Craig JM et al. Patients with familial adenomatous polyposis harbor colonic biofilms containing tumorigenic bacteria. Science 2018; 359: 592–597.

31 Bracht K, Nicholls AM, Liu Y, Bodmer WF. 5-Fluorouracil response in a large panel of colorectal cancer cell lines is associated with mismatch repair deficiency. Br J Cancer 2010; 103: 340–346.

32 Spanogiannopoulos P, Bradley PH, Melamed J et al. Drug resistant gut bacteria mimic a host mechanism for anticancer drug clearance. bioRxiv 2019;: 820084.

33 Davidson AL, Chen J. ATP-Binding Cassette Transporters in Bacteria. Annu Rev Biochem 2004; 73: 241–268.

34 King AE, Ackley MA, Cass CE, Young JD, Baldwin SA. Nucleoside transporters: from scavengers to novel therapeutic targets. Trends Pharmacol Sci 2006; 27: 416–425.

35 Acimovic Y, Coe IR. Molecular evolution of the equilibrative nucleoside transporter family: identification of novel family members in prokaryotes and eukaryotes. Mol Biol Evol 2002; 19:2199–2210.

36 Kumari S, Tripathi P. Nucleotide metabolism pathway: the achilles’ heel for bacterial pathogens. Curr Sci 2021; 120: 6.

37 Vogels GD, Van der Drift C. Degradation of purines and pyrimidines by microorganisms. Bacteriol Rev 1976; 40: 403–468.

38 Yamamura K, Baba Y, Nakagawa S et al. Human Microbiome Fusobacterium Nucleatum in Esophageal Cancer Tissue Is Associated with Prognosis. Clin Cancer Res Off J Am Assoc Cancer Res 2016; 22: 5574–5581.

39 García-González AP, Ritter AD, Shrestha S, Andersen EC, Yilmaz LS, Walhout AJM. Bacterial Metabolism Affects the C. elegans Response to Cancer Chemotherapeutics. Cell 2017; 169: 431–441.e8.

40 Scott TA, Quintaneiro LM, Norvaisas P et al. Host-Microbe Co-metabolism Dictates Cancer Drug Efficacy in C. elegans. Cell 2017; 169: 442–456.e18.

41 Wei X, McLeod HL, McMurrough J, Gonzalez FJ, Fernandez-Salguero P. Molecular basis of the human dihydropyrimidine dehydrogenase deficiency and 5-fluorouracil toxicity. J Clin Invest 1996; 98: 610–615.

42 van Kuilenburg ABP. Dihydropyrimidine dehydrogenase and the efficacy and toxicity of 5-fluorouracil. Eur J Cancer Oxf Engl 1990 2004; 40: 939–950.

43 Hidese R, Mihara H, Kurihara T, Esaki N. Escherichia coli dihydropyrimidine dehydrogenase is a novel NAD-dependent heterotetramer essential for the production of 5,6-dihydrouracil. J Bacteriol 2011; 193: 989–993.

44 Alexander JL, Wilson ID, Teare J, Marchesi JR, Nicholson JK, Kinross JM. Gut microbiota modulation of chemotherapy efficacy and toxicity. Nat Rev Gastroenterol Hepatol 2017; 14: 356–365.

45 Geller LT, Barzily-Rokni M, Danino T et al. Potential role of intratumor bacteria in mediating tumor resistance to the chemotherapeutic drug gemcitabine. Science 2017; 357: 1156–1160.

46 Saam J, Critchfield GC, Hamilton SA, Roa BB, Wenstrup RJ, Kaldate RR. Body surface area-based dosing of 5-fluoruracil results in extensive interindividual variability in 5-fluorouracil exposure in colorectal cancer patients on FOLFOX regimens. Clin Colorectal Cancer 2011; 10: 203–206.

47 Zhang N, Yin Y, Xu S-J, Chen W-S. 5-Fluorouracil: mechanisms of resistance and reversal strategies. Mol Basel Switz 2008; 13: 1551–1569.

48 Pratt S, Shepard RL, Kandasamy RA, Johnston PA, Perry W, Dantzig AH. The multidrug resistance protein 5 (ABCC5) confers resistance to 5-fluorouracil and transports its monophosphorylated metabolites. Mol Cancer Ther 2005; 4: 855–863.

49 Higgins CF. ABC transporters: from microorganisms to man. Annu Rev Cell Biol 1992; 8: 67–113.

50 Rosener B, Sayin S, Oluoch PO et al. Evolved bacterial resistance against fluoropyrimidines can lower chemotherapy impact in the Caenorhabditis elegans host. eLife 2020; 9: e59831.

51 Strauss J, Kaplan GG, Beck PL et al. Invasive potential of gut mucosa-derived Fusobacterium nucleatum positively correlates with IBD status of the host. Inflamm Bowel Dis 2011; 17: 1971–1978.

